# A gut pathobiont synergizes with the microbiota to instigate inflammatory disease marked by immunoreactivity against other symbionts but not itself

**DOI:** 10.1101/178038

**Authors:** João Carlos Gomes-Neto, Hatem Kittana, Sara Mantz, Rafael R. Segura Munoz, Robert J. Schmaltz, Laure B. Bindels, Jennifer Clarke, Jesse M. Hostetter, Andrew K. Benson, Jens Walter, Amanda E. Ramer-Tait

## Abstract

Inflammatory bowel diseases (IBD) are likely driven by aberrant immune responses directed against the resident microbiota. Although IBD is commonly associated with a dysbiotic microbiota enriched in putative pathobionts, the etiological agents of IBD remain unknown. Using a pathobiont-induced intestinal inflammation model and a defined bacterial community, we provide new insights into the immune-microbiota interactions during disease. In our model system, the pathobiont *Helicobacter bilis* instigates disease following sub-pathological dextran sulfate sodium treatment. We show that *H. bilis* causes mild inflammation in mono-associated mice, but severe disease in the presence of a microbiota, demonstrating synergy between the pathobiont and microbiota in exacerbating pathology. Remarkably, inflammation depends on the presence of *H. bilis*, but is marked by a predominant Th17 response against specific members of the microbiota and not the pathobiont, even upon the removal of the most immune-dominant taxa. Neither increases in pathobiont burden nor unique changes in immune-targeted microbiota member abundances are observed during disease. Collectively, our findings demonstrate that a pathobiont instigates inflammation without being the primary target of a Th17 response or by altering the microbiota community structure. Moreover, our findings point toward monitoring pathobiont-induced changes in microbiota immune targeting as a new concept in IBD diagnotics.

## Introduction

IBD is associated with an altered gut microbiota composition (i.e., dysbiosis) characterized by a loss of members belonging to the Firmicutes and Bacteroidetes phyla and an expansion of Proteobacteria ^1^. Although cause-and-effect-relationships for disease-related dysbioses have not been established ^2-4^, the microbiota associated with IBD has been hypothesized to be enriched in pathobionts—microorganisms that are thought to exert a pathological role when their relationship with the host is altered ^5,6^. Unlike frank pathogens, pathobionts presumably live symbiotic lifestyles under normal circumstances without negatively affecting host health; however, they are capable of selectively expanding during episodes of inflammation and may exacerbate the disease process ^7,8^. It is believed that pathobionts can instigate and/or perpetuate pro-inflammatory responses; however, the exact immune-microbiota interactions governing pathobiont-mediated intestinal pathology are not well understood ^9-17^.

Here, we explore these inter-relationships using a mouse model harboring the Altered Schaedler Flora (ASF) in which disease can be induced through the addition of the pathobiont *Helicobacter bilis* and a sub-pathological dose of dextran sulfate sodium (DSS) ^18^. Although *H. bilis* and other enterohepatic Helicobacters are well-known for contributing to intestinal inflammation in immunocompromised or inflamed hosts ^18-22^, the role of particular microbiota members in mediating pathology has not been elucidated due to the difficulty in determining specific immune-microbiome interactions in conventional animals. By using the defined ASF community in both wild-type and *Rag1*^-/-^ mice, we systematically identified (i) the relative contribution of the pathobiont and the microbiota to disease severity, (ii) to what degree disease was associated with changes in the microbial community composition, (iii) the role of the adaptive immune system in disease, (iv) the specific microbes towards which immune responses were directed and (v) the contributions of immune-targeted taxa to the disease process.

## Results

### Pathobiont colonization can induce an equivalent degree of disease in mice with a conventional or defined microbiota

To study specific immune-microbe interactions in pathobiont-mediated colitis, we first established that a mouse model with a defined microbial community can replicate the severity of pathobiont-induced intestinal inflammation observed in the same mouse line harboring a conventional (e.g., complex and undefined) microbiota. To this end, C3H/HeN adult mice (8-10 weeks old) carrying either a conventional (Conv) microbiota or the ASF community from birth were colonized with *H. bilis* for three weeks prior to treatment with a sub-pathological dose of DSS (1.5%). Disease severity was not significantly different between Conv and ASF mice (Fig. 1A), and no differences in *H. bilis* abundance were observed across treatments (Fig. 1B). In addition, *H. bilis*-DSS-mediated inflammation did not result in major alterations to the abundances of individual bacteria or the community structure in ASF mice, except for decreased numbers of ASF 492, which was decreased in all mice treated with DSS (Fig. 1C-D). Homogeneity of variances analysis (Betadisper) using Bray-Curtis dissimilarity coefficients further confirmed the absence of differences in ASF community dispersion across treatments (*P* = 0.573) as did results from PERMANOVA (R^2^ = 21.28, *P* = 0.003) and ANOSIM (R = 0.133) analyses, which found only minor treatment effects. Because *H. bilis* colonization exacerbated disease equivalently in both Conv and ASF microbiota mice following DSS treatment, this defined model now permits the study of exact immune-microbe interactions during disease.

**Figure 1.**
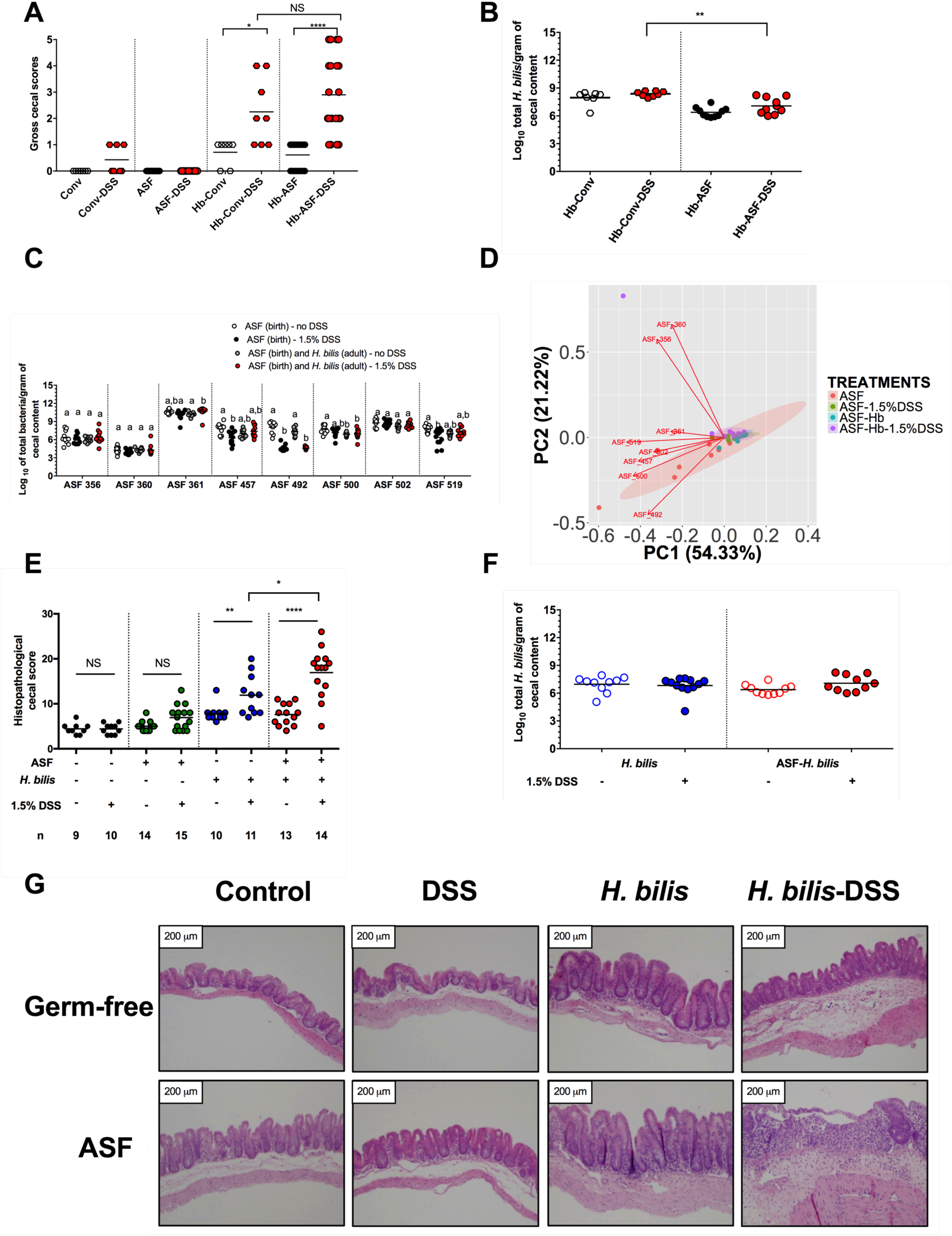
Colonization with the pathobiont *Helicobacter bilis* induced severe intestinal inflammation in the presence of a microbiota but did not alter microbiota composition. (**A**) Gross cecal scores depicting disease severity for Conventional (Conv) and Altered Schaedler Flora (ASF) mice colonized with or without the pathobiont *H. bilis* (Hb) for 3 wks and then either treated with 1.5% DSS or left untreated (*n* = 7-22 animals per treatment). (**B**) *H. bilis* abundance in cecal contents from Conv and ASF mice colonized with Hb for 3 wks and then treated with or without DSS (*n* = 7-10 animals per treatment). (**C**) ASF bacterial abundances in cecal contents of ASF-bearing animals colonized with Hb and either treated with 1.5% DSS or left untreated at 11-12 wks of age (*n* = 10-12 animals per treatment). (**D**) Principal Component Analysis (PCA) of the ASF community structure for mice from all treatments. The PCA plot dots represent the entire ASF community based on abundance of each organism as measured by species-specific qPCR assays in cecal contents (*n* = 10-12 animals per treatment). The red arrows in the PCA plot depict the directionality and magnitude of each ASF member’s contribution to the variability across both PC1 and PC2. (**E**) Histopathological disease scores for cecal tissues from germ-free (GF), ASF-bearing, *H. bilis* mono-associated and *H. bilis* colonized ASF-bearing mice either treated with 1.5% DSS or left untreated. The letter *n* in the graph indicates the number of mice used per treatment group. Scores were based on presence or absence of morphological alterations such as stromal collapse or ulceration, gland hyperplasia, inflammatory cell infiltrate in the lamina propria and submucosal edema across groups. (**F**) Pathobiont abundance measured by qPCR in cecal contents of mono-associated or ASF-bearing mice treated with DSS or left untreated (*n* = 10-12 animals per treatment, each represented by an individual circle). (**G**) Representative photomicrographs of hematoxylin-eosin (H&E) stained tissues were taken at 100X magnification. Images were selected based on two criteria: (1) representing the mean histopathological score for each group and (2) by considering the pathologist’s description of the morphological changes. Horizontal bars represent group means in all graphs. (**C**) Differing superscript letters indicate significant differences across treatments as per a non-parametric Kruskal-Wallis one-way ANOVA, followed by a post-hoc test (Dunn’s test, *P* < 0.05). (**A**-**B** and **E-F**) Asterisks refer to the degree of significance for differences as determined by a non-parametric unpaired Mann-Whitney test using a two-tailed distribution for *P*-value calculations (* *P* < 0.05, ** *P* ≤ 0.01, *** *P* ≤ 0.001, **** *P* ≤ 0.0001 and NS = not significant at *P* ≥ 0.05). No comparisons in Fig. 1F were significant (*P* ≥ 0.05). Only significant differences between treatments are presented in the graphs with the exception of one comparison in Fig. 1A. Pathobiont and ASF bacterial abundances were measured using species-specific qPCR assays. Experiments were performed using male and female C3H/HeN mice at 8-10 wks of age; mice were colonized with *H. bilis* for 3 wks.

### The resident microbiota acts synergistically with the pathobiont to induce severe disease

Having established that the pathobiont plus the ASF community are sufficient to cause disease following low-dose DSS treatment, we next determined the relative contributions of the resident microbiota and pathobiont to the pathology. Groups of adult germ-free and ASF-bearing C3H/HeN mice were colonized with or without *H. bilis* for three weeks and then either treated with low-dose DSS or left untreated. Histopathological evaluation of cecal tissues revealed that ASF-bearing mice colonized with *H. bilis* and treated with DSS developed severe intestinal lesions characterized by stromal collapse and epithelial ulceration, increased inflammatory infiltrate, submucosal edema and gland hyperplasia (Fig. 1E and G; Fig. S1A). In contrast, only moderate lesions were observed in mice mono-associated with *H. bilis* and treated with DSS. Of note, histopathological analysis demonstrated that germ-free and ASF control mice treated with 1.5% DSS developed only mild microscopic lesions, confirming that this low dose itself is sub-pathological (Fig. 1E) as previously reported ^18^. Enumeration of *H. bilis* by qPCR revealed no significant effects of the ASF community and/or DSS treatment on pathobiont abundance (Fig. 1F). Thus, even though *H. bilis* by itself induces moderate disease following a low-dose of DSS, the presence of the ASF community is necessary for severe disease, indicating that the resident microbiota exacerbates pathobiont-mediated inflammation.

### The pathobiont and resident microbiota synergistically induce a pro-inflammatory response associated with disease

Analysis of cytokine profiles from the cecal tissues of *H. bilis*-ASF-DSS mice showed that severe intestinal inflammation is marked by a significant increase in the production of several cytokines considered hallmarks of a Th17 response. This response included elevation of TGF-ß1 and the pro-inflammatory cytokines IL-1ß, IL-17A and IL-17F (Fig. 2A-D) as well as a corresponding decrease in production of the anti-inflammatory cytokine IL-10 (Fig. 2E) as compared to either *H. bilis*-DSS or ASF-DSS mice. IL-33 and IFN-γ production were also increased in the inflamed ceca of *H. bilis*-ASF-DSS mice (Fig. S1B-C). Production of TNF-α and IL-22 was elevated in *H. bilis*-ASF versus *H. bilis* mono-associated mice, but DSS treatment did not further increase the secretion of either cytokine (Fig. S1D-E). Evaluation of immune cell populations via flow cytometry revealed an expansion of effector memory (EM) CD44^high^ CD62L^low^ CD4^+^ T cells (Fig. 2F) in *H. bilis*-ASF-DSS mice compared to either ASF-DSS or *H. bilis*-ASF mice. Together, the Th17 signature observed in the pro-inflammatory cytokine profile and the expansion of activated CD4^+^ T cells imply a contributing role for adaptive immunity in precipitating severe intestinal inflammation and pathology in our model.

**Figure 2.**
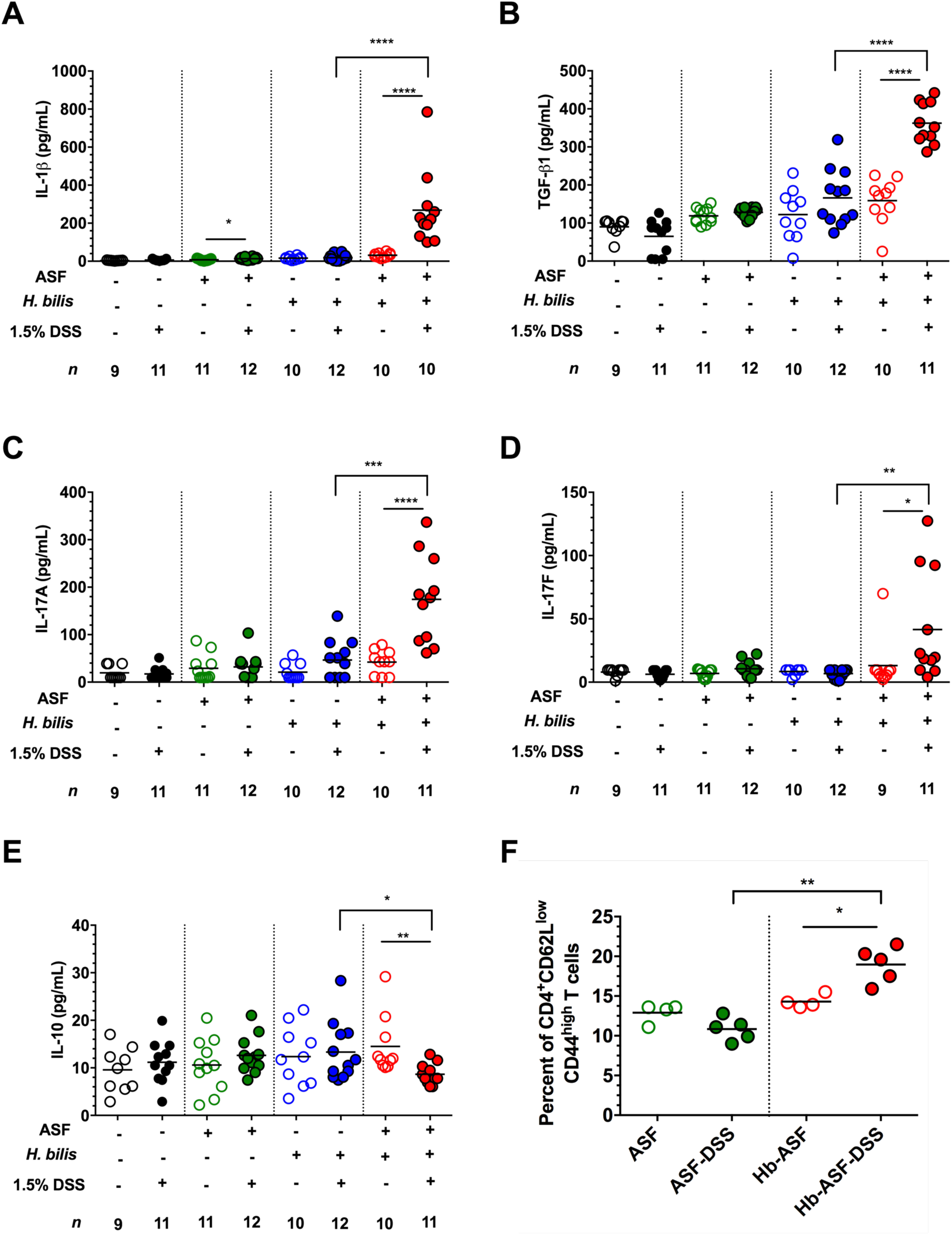
The resident microbiota acted synergistically with the pathobiont to induce a pro-inflammatory immune response. (**A-E**) Pro- and anti-inflammatory cytokines present in cecal explants from germ-free (GF), ASF-bearing, *H. bilis* mono-associated and *H. bilis* colonized ASF bearing mice treated with 1.5% DSS or left untreated (*n* = 9-12 animals per treatment). (**F**) Proportion of effector memory (EM) CD44^high^CD62L^low^ CD4^+^ T cells in mesenteric lymph nodes (each dot represents a pool of 2-3 animals) of ASF-bearing mice colonized with or without *H. bilis* and either treated with DSS or left untreated. (**A-F**) Horizontal bars represent treatment means in all graphs. Asterisks depict the degree of significance for differences as determined by a non-parametric unpaired Mann-Whitney test using a two-tailed distribution for *P*-value calculations (* *P* < 0.05, ** *P* ≤ 0.01, *** *P* ≤ 0.001 and **** *P* ≤ 0.0001). Only significant differences between treatments are presented in the graphs. Experiments were performed using male and female C3H/HeN mice at 8-10 wks of age; mice were colonized with *H. bilis* for 3 wks.

### The adaptive immune system is required for severe pathobiont-mediated intestinal inflammation

To assess the role of the adaptive immune response in pathobiont mediated-disease, we switched our mouse model to the C57BL/6 background to compare disease severity for the *H. bilis*-ASF-DSS treatment in isogenic wild-type (WT) and *Rag1*^*-/-*^ mice, the latter of which lack mature T and B cells. In this genetic background, WT *H. bilis*-ASF-DSS mice develop disease that is equivalent in severity to that observed in the C3H/HeN line. In contrast, *Rag1*^*-/-*^ *H. bilis*-ASF-DSS mice experienced significantly less severe disease compared to their WT counterparts, confirming a pathological role for the adaptive immune system in pathobiont-mediated disease (Fig. 3A). Pathobiont abundance was not significantly altered by adaptive immune status alone (Fig. 3B). The *Rag1*^*-/-*^-dependent reduction of disease was characterized by a marked decrease in local cytokine and chemokine production, including IL-1ß and IL-17 (Fig. 3C-D) as well as G-CSF, MIP-2 and IL-6 (Fig. S2A-C). In contrast, GM-CSF, MCP-1, IL-1α, IL-33 and IFN-γ were not affected by the *Rag1* deletion and remained high in both WT and *Rag1*^*-/-*^ *H. bilis*-ASF-DSS mice (Fig. S2D-H). The severe disease observed in WT C57BL/6 *H. bilis*-ASF-DSS mice was accompanied by a significant expansion of both EM CD4^+^ T cells and Th17 cells (Fig. 3E-F and Fig. S3A). Increased populations of activated B cells and iTregs (CD25^+^ Foxp3^+^ Neuropilin-1^low^ Helios^low^ CD4^+^) were also observed in WT C57BL/6 mice with pathobiont-induced disease (Fig. S3B-C). Total numbers of IFN-γ^+^ CD4^+^ T cells, IL-17A^+^ IFN-γ^+^ CD4^+^ T cells, Tregs (CD25^+^ Foxp3^+^ CD4^+^), nTregs (CD25^+^ Foxp3^+^ Neuropilin-1^high^ Helios^high^ CD4^+^) and CD8b^+^ T cells remained unaltered (Fig. S3D-J).

**Figure 3.**
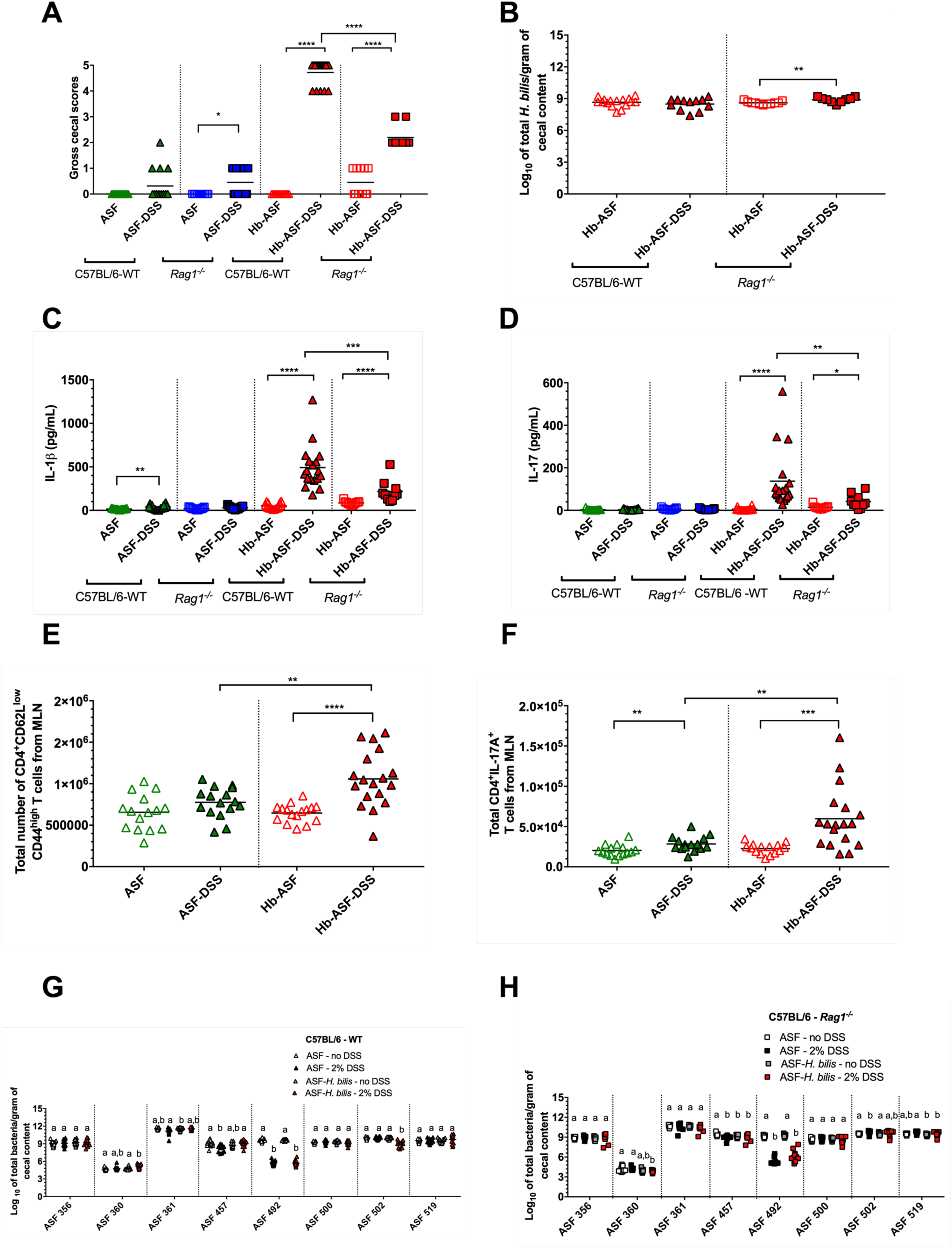
The absence of adaptive immunity limited the severity of pathobiont-induced intestinal inflammation and the magnitude of the Th17 response without altering microbiota composition. (**A**) Gross cecal scores depicting disease in ASF-bearing C57BL/6 wild type (WT) and *Rag1*^*-/-*^ mice harboring the ASF and colonized with or without *H. bilis* (Hb) and either treated with 2% DSS or left untreated (*n* = 10-18 animals per treatment). (**B**) Pathobiont abundance in cecal contents as measured by species-specific qPCR (*n* = 9-16 animals per treatment). (**C-D**) IL-1ß (**C**) and IL-17 (**D**) concentrations in cecal explants (*n* = 10-18 animals per treatment). (**E-H**) Total numbers of effector memory (EM) CD44^high^ CD62L^low^ CD4^+^ T cells (E) and total IL-17A^+^ CD4^+^ T cells (**F**) in mesenteric lymph nodes (MLN; *n* = 15-18 animals per treatment). (**G-H**) ASF bacterial abundances in cecal contents of ASF-bearing C57BL/6 WT and *Rag1*^*-/-*^ mice harboring the ASF and colonized with or without *H. bilis* and either treated with 2% DSS or left untreated (*n* = 9-18 animals per treatment). (**A-F**) Horizontal bars represent treatment means in all graphs. Differences in the absolute immune cell counts were tested using an unpaired parametric T-test. A non-parametric unpaired Mann-Whitney test was used to evaluate the gross cecal scores, pathobiont abundance and cytokine concentrations. All *P*-values were calculated, across all tests, using a two-tailed distribution. Asterisks show the degree of significance for the indicated comparisons in each graph (\**P* < 0.05, ** *P* ≤ 0.01, *** *P* ≤ 0.001 and **** *P* ≤ 0.0001). Only significant differences between treatments are presented in the graphs. (**G-H**) Differing superscript letters indicate significant differences across treatments as per a non-parametric Kruskal-Wallis one-way ANOVA, followed by a post-hoc test (Dunn’s test, *P* < 0.05). ASF bacterial abundances were measured using species-specific qPCR assays. Experiments were performed using male and female WT or *Rag1*^-/-^ C57BL/6 mice at 8-10 wks of age; mice were colonized with *H. bilis* for 3 wks.

Analysis of ASF bacterial abundances in the C57BL/6 mouse line showed no major differences between either WT or *Rag1*^-/-^ mice colonized with *H. bilis* and treated with DSS (Fig. 3G-H). As observed in previous experiments, DSS treatment significantly decreased the abundance of ASF 492 (*E. plexicaudatum*) independently of pathobiont status. The absence of significant changes in microbiota structure were further confirmed using Betadisper (*P* = 0.098 and *P* = 0.17 for WT and *Rag1*^-/-^, respectively), ANOSIM (R statistics = 0.1331 and 0.1224 for WT and *Rag1*^-/-^, respectively) and PERMANOVA (nearly 15% of both WT and *Rag1*^-/-^ dataset variances were explained by treatments) analyses.

### Pathobiont colonization triggers CD4^+^ Th17 cell immunoreactivity against specific members of the resident microbiota, but not toward itself

Our finding that severe *H. bilis*-mediated disease requires a microbiota and is associated with IL-17 production led us to next determine which of the pathobiont and defined ASF bacterial community members are the actual targets of this Th17 responses. CD4^+^ T cells were isolated from the mesenteric lymph nodes of either germ-free or ASF-bearing C3H/HeN mice with or without *H. bilis* colonization for three weeks prior to low-dose DSS treatment as described in Fig. 1E. Cytokine profiles were then assessed in supernatants of these CD4^+^ T cells after co-culturing with mitomycin C-treated naïve splenocytes pulsed with antigens derived from either *H. bilis* or individual ASF members or with unpulsed splenocytes. Cytokine production was considered specific to a certain ASF member if there were significant increases over (i) unstimulated T cells isolated from the same mice, (ii) antigen-stimulated T cells harvested from equivalently-treated mice not harboring the ASF community and (iii) antigen-stimulated T cells from ASF-bearing mice treated with DSS but not colonized with *H. bilis*.

CD4^+^ T cells isolated from severely diseased *H. bilis*-ASF-DSS mice produced significantly elevated levels of IL-17A when stimulated with antigens from either ASF 356, ASF 361 and, to a lesser extent, ASF 502 as compared to *H. bilis* mono-associated-DSS mice (Fig. 4A-B, 4E-F; Fig. S4D). Antigen-specific responses to ASF 360, 492, 500 and 519 were negligible (Fig. S4A-C and E) and those to 457 were undetectable (Fig. 4C). Interestingly, IL-17A immunoreactivity specifically against the *H. bilis* pathobiont itself was not observed (Fig. 4D). Importantly, qPCR analysis of cecal tissues and contents showed no correlation between the abundances of the pathobiont and species primarily targeted by the immune system (i.e., ASF 356 and ASF 361) during disease (Fig. S4F-H).

**Figure 4.**
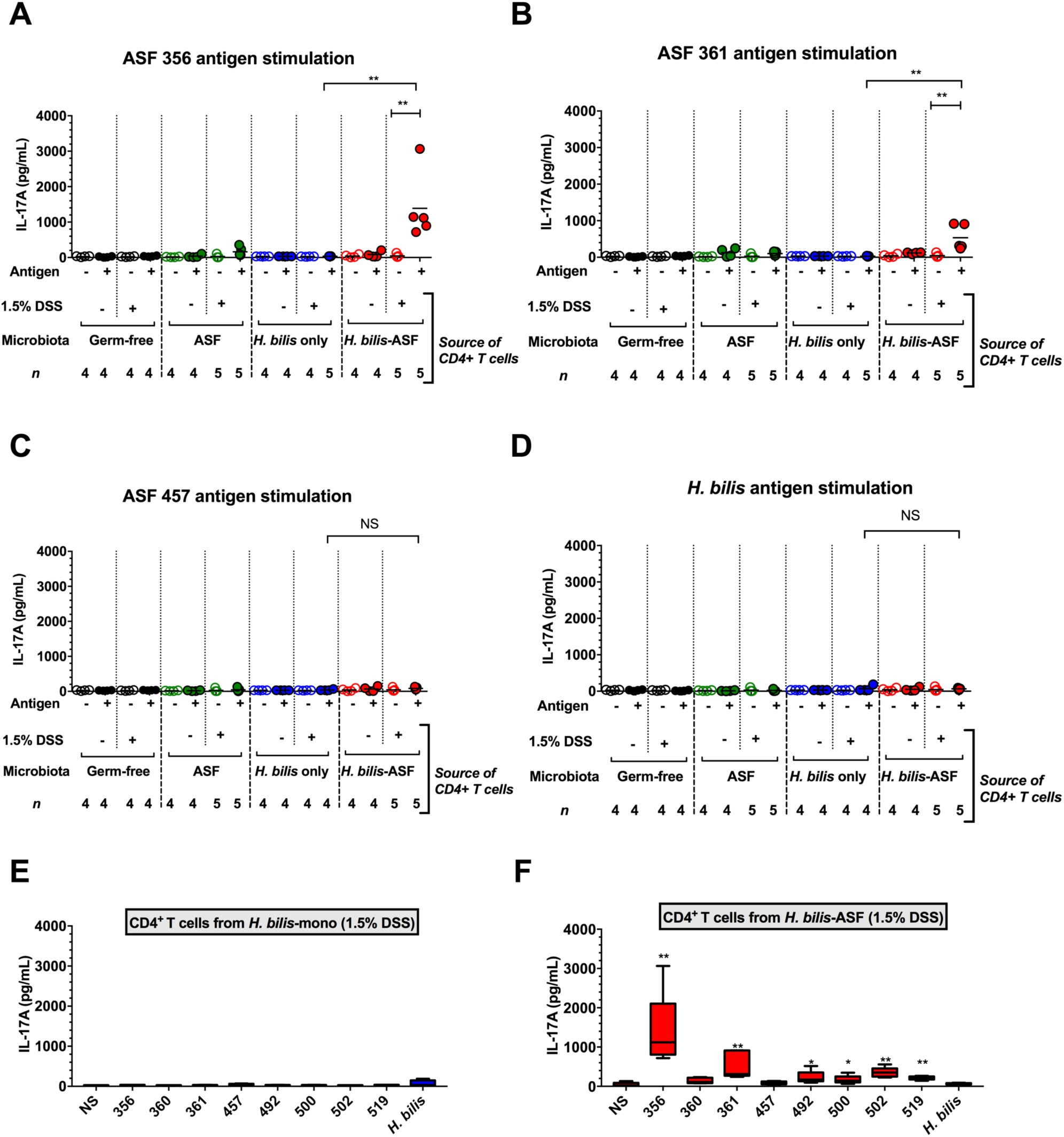
Pathobiont colonization triggered CD4^+^ Th17 cell immunoreactivity against specific members of the resident microbiota but not toward itself. (**A-B**) IL-17A secretion from CD4^+^ T cells isolated from the mesenteric lymph nodes of germ-free (GF), ASF-bearing, *H. bilis* mono-associated and *H. bilis* colonized ASF bearing mice treated with 1.5% DSS or left untreated. CD4^+^ T cells were either left unstimulated (NS) or stimulated with whole-cell sonicate antigens from either ASF 356 (*Clostridium sp.*) (**A**), ASF 361 (*Lactobacillus murinus*) (**B**), ASF 457 (*Mucispirillum schaedleri*) (**C**) or the pathobiont (*H. bilis*) (**D**) for 72 hrs. (**E-F**) IL-17A secretion from CD4^+^ T cells isolated from the mesenteric lymph nodes of either *H. bilis* mono-associated mice (**E**) or ASF-bearing mice colonized with *H. bilis* (**F**) and stimulated with whole-cell sonicate antigens from each of the individual ASF members or *H. bilis* (*n* = 4-5 pools of 2-3 animals per pool per treatment). Pairwise comparisons in (**E**) and (**F**) were performed with respect to unstimulated cells. Box-and-whiskers plot depict minimum and maximum values in addition to the median (i.e., horizontal bar in the middle). (**A-D**) Horizontal bars represent treatment means in all graphs. (**A-F**) Asterisks depict the degree of significance for differences as determined by a non-parametric unpaired Mann-Whitney test using a two-tailed distribution for *P*-value calculations (* *P* < 0.05, ** *P* ≤ 0.01, *** *P* ≤ 0.001, **** *P* ≤ 0.0001 and NS = not significant *P* ≥ 0.05). Experiments were performed using male and female C3H/HeN mice at 8-10 wks of age; mice were colonized with *H. bilis* for 3 wks.

Similar to IL-17A, the same patterns of immunoreactivity to ASF 356 and 361, but not the pathobiont, were also observed for IL-17F (Fig. S5A-I). Antigen-specific IL-22 production from CD4^+^ T cells harvested from *H. bilis*-ASF-DSS mice was observed upon stimulation with antigens from ASF 356 but not following stimulation with *H. bilis* antigens (Fig. S6A-I). Although ASF antigen-specific IFN-γ and IL-10 responses were observed, they were not unique to the severely diseased *H. bilis*-ASF-DSS mice (Fig. S7A-I and S8A-I). Given the requirement of both the pathobiont and ASF microbiota for severe disease, the pattern of immunoreactivity targeting immune-dominant ASF members but not *H. bilis* demonstrates that this pathobiont can elicit Th17 immune responses against the resident microbiota and bring about intestinal inflammation without being the primary immunological target.

### Removing immune-dominant gut symbionts redirects the Th17 immune response to other ASF members and does not alter the severity of pathobiont-mediated disease

To determine the role of the immune-dominant members ASF 356 and ASF 361 in pathobiont-induced disease, we next compared disease susceptibility in *H. bilis* colonized and DSS-treated mice harboring either a complete ASF community or ASF communities lacking immune dominant strains (Fig. 5A). Specifically, prior to low-dose DSS exposure, we colonized germ-free mice with either (i) *H. bilis* plus all eight ASF members, (ii) *H. bilis* plus the ASF without 356 (the major immune-dominant member), (iii) *H. bilis* plus the ASF without 356 and 361 (the two most immune-dominant members), and (iv) *H. bilis* plus the ASF without 457 (an ASF member that is not immune targeted). Surprisingly, removal of the immune-dominant ASF members 356 and/or 361 did not limit disease severity (Fig. 5B), and the pathobiont abundance remained unaltered (Fig. 5C). Interestingly, ASF 361 became the primary target of the Th17 response in the absence of ASF 356 (Fig. 5D-F). Similarly, when both ASF 356 and 361 were not present, ASF 502 became the immune dominant symbiont with other members also being targeted, albeit to lesser degree (Fig. 5D-F; Fig. S9A-L). Also notable was the observation that even in these communities of lower complexity, Th17 immunoreactivity against both ASF 457 and *H. bilis* remained negligible (Fig. 5G-H; Fig. S9H and M).

**Figure 5.**
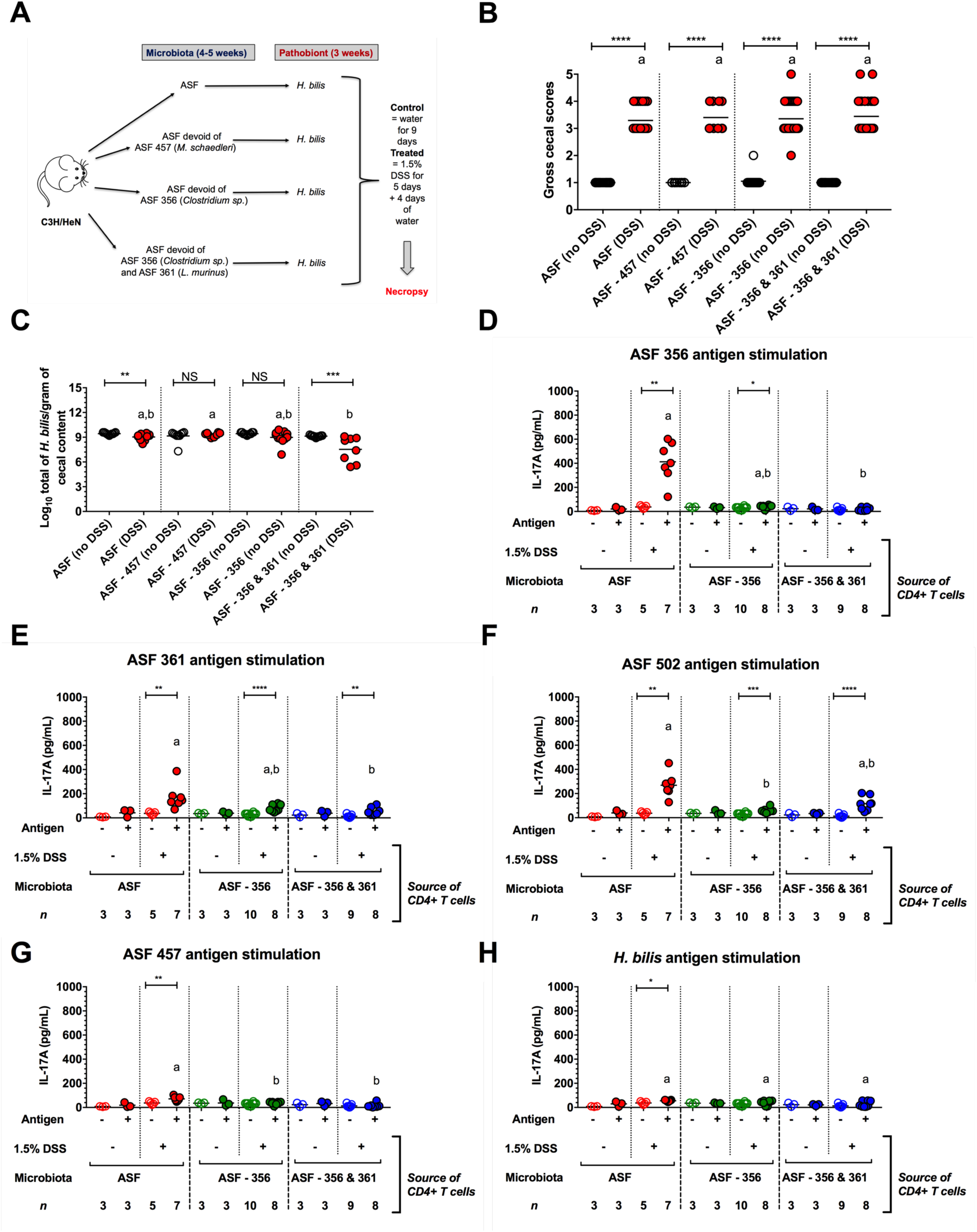
Removal of immune-dominant gut symbionts did not alter the severity of pathobiont-mediated disease and precipitated a shift in the pattern of Th17 immunoreactivity. (**A**) Experimental design describing all treatment groups which included all ASF members or ASF communities devoid of the following taxa: ASF 356 (*Clostridium sp.*), ASF 356 and 361 (*Lactobacillus murinus*), or ASF 457 (*Mucispirillum schaedleri*). Mice harboring these communities were then colonized with *H. bilis* for 3 wks and then either treated with 1.5% DSS or left untreated at 12-13 wks of age. (**B**) Gross cecal scores depicting disease severity (*n* = 10-28 animals per treatment). (**C**) *H. bilis* abundance in cecal contents as measured by species-specific qPCR (*n* = 7-10 animals per treatment). (**D-H**) IL-17A secretion from CD4^+^ T cells isolated from the mesenteric lymph nodes of mice in all treatments except for those colonized with the ASF minus 457. CD4^+^ T cells were either left unstimulated (NS) or stimulated with whole-cell sonicate antigens from either ASF 356 (*Clostridium sp.*) (**D**), ASF 361 (*L. murinus*) (**E**), ASF 502 (*Clostridium sp.*) (**F**), ASF 457 (*M. schaedleri*) (**G**) or the pathobiont (*H. bilis*) (**H**) for 72 hrs (*n* = 3-10 pools of 2-3 animals per pool per treatment). (**B-H**) Asterisks depict the degree of significance for differences as determined by a non-parametric unpaired Mann-Whitney test using a two-tailed distribution for *P*-value calculations (* *P* < 0.05, ** *P* ≤ 0.01, *** *P* ≤ 0.001, **** *P* ≤ 0.0001 and NS = not significant *P* ≥ 0.05). (**B-H**) Differing superscript letters indicate significant differences across treatments as per a non-parametric Kruskal-Wallis one-way ANOVA, followed by a post-hoc test (Dunn’s test, *P* < 0.05). Experiments were initiated when male and female C3H/HeN mice were 4-5 wks of age.

Some changes in individual ASF member abundances were observed following removal of the immune-dominant species, including increased numbers of ASF 360 (Fig. 6B), an expansion of ASF 361 upon removal of ASF 356 or ASF 457 (Fig. 6C-D), and, as seen in previous experiments, decreased numbers of ASF 492 following DSS treatment regardless of the community composition (Fig. 6E). However, no patterns of microbial community structure were observed that were unique to pathobiont-induced disease. Of note, the immune-dominant ASF members targeted by the Th17 response were not enriched in the IgA^+^ fraction of cecal contents during disease (Fig. S10A-K). Collectively, these findings demonstrate that removal of the immune-dominant gut symbiont(s) during pathobiont-mediated inflammation maintains pathobiont-mediated Th17 immune responses against remaining members and not the pathobiont itself without affecting the disease severity.

**Figure 6.**
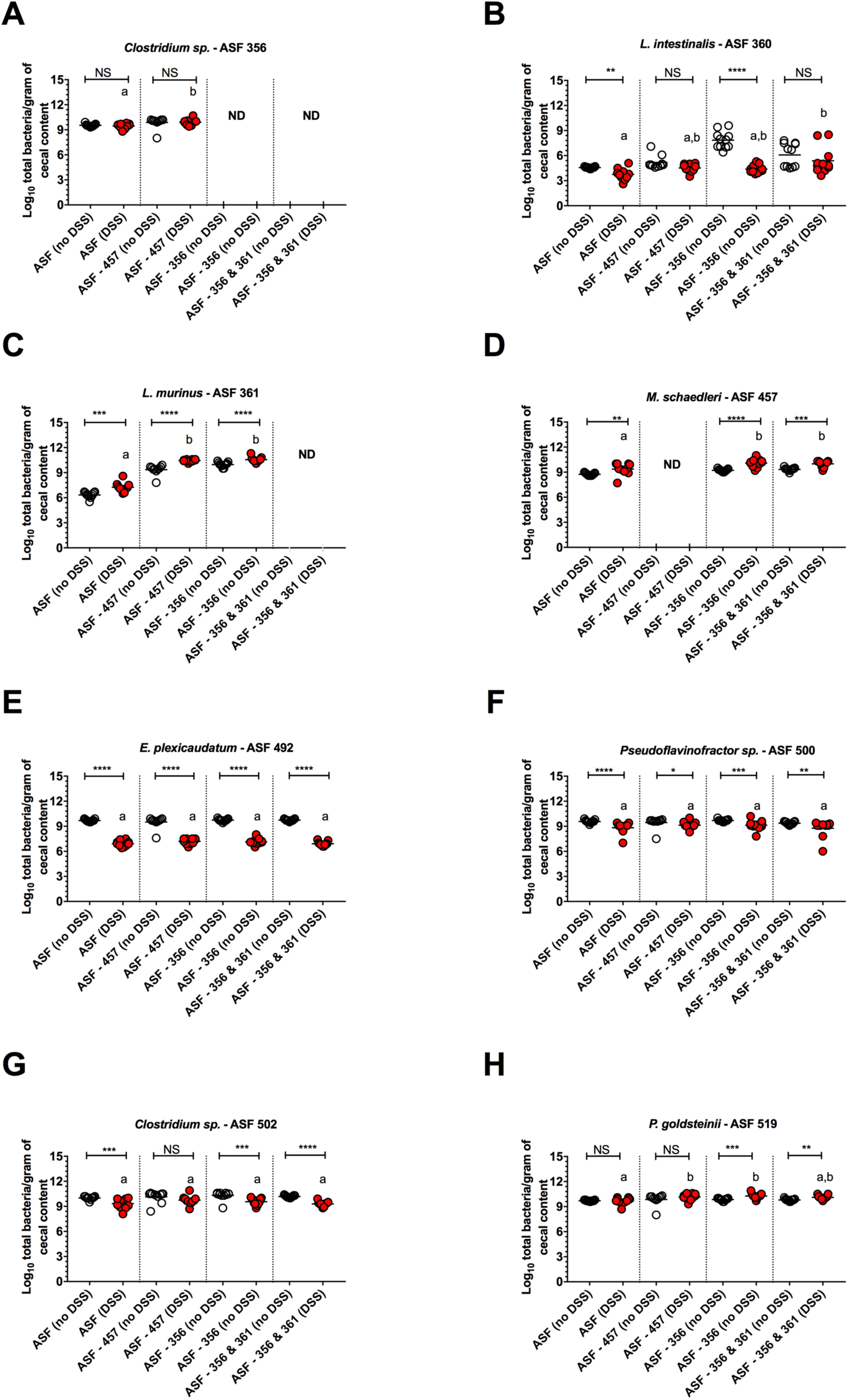
Removal of immune-dominant gut symbionts did not create patterns of microbial community structure that were unique to pathobiont-induced disease. (**A-H**) ASF bacterial abundances in cecal contents of animals colonized with all ASF members or devoid of the following taxa: ASF 356 (*Clostridium sp.*), 356 and 361 (*Lactobacillus murinus*), or 457 (*Mucispirillum schaedleri*). Mice harboring these communities were then colonized with *H. bilis* for 3 wks and then either treated with 1.5% DSS or left untreated at 12-13 wks of age (*n* = 10-11 animals per treatment). Asterisks depict the degree of significance for differences as determined by a non-parametric unpaired Mann-Whitney test using a two-tailed distribution for *P*-value calculations (* *P* < 0.05, ** *P* ≤ 0.01, *** *P* ≤ 0.001, **** *P* ≤ 0.0001 and NS = not significant *P* ≥ 0.05). Differing superscript letters indicate significant differences across treatments as per a non-parametric Kruskal-Wallis one-way ANOVA, followed by a post-hoc test (Dunn’s test, *P* < 0.05). ND refers to not detectable. ASF bacterial abundances were measured using species-specific qPCR assays. Experiments were initiated when male and female C3H/HeN mice were 4-5 wks of age.

## Discussion

IBD is hypothesized to result from aberrant immune responses directed against gut symbionts, which can be initiated by the loss of intestinal barrier integrity ^23-25^. These chronic diseases are routinely characterized by microbial dysbiosis, including overrepresentation of the phylum Proteobacteria ^3,4,26-28^. Certain Proteobacteria are thought to behave as pathobionts during IBD, and much like the different virulence strategies found among frank pathogens belonging to this phylum, multiple strategies seem to have evolved in pathobionts that enable them to contribute to disease. These general strategies include altering barrier function, invading the gut epithelium, and/or stimulating local inflammatory responses, but the mechanisms through which these strategies manifest are not well known ^5,6^. To define specific mechanisms that underlie general pathobiont strategies, we focused on the pathobiont *H. bilis*, which is capable of triggering inflammatory disease in immunocompromised rodent models ^18,20-22,29^. In our current model of immunosufficient, gnotobiotic mice colonized with *H. bilis*, a subpathological perturbation of the intestinal barrier (DSS treatment) is required for disease, but is not sufficient because severe pathology also requires a microbiota. Using a simplified microbiota (ASF), we show that *H. bilis* precipitates severe disease without significant increases in its own abundance and without substantial changes in gut microbial community structure. These features differentiate the pathobiont characteristics of *H. bilis* from other pathobionts such as the Adherent and Invasive *Escherichia coli* strain LF82 (AIEC LF82), which elicits chronic colitis and brings about changes in microbiota composition after disruption of innate immune signaling ^11,30^.

Perhaps the most striking characteristic of the severe intestinal inflammation observed in ASF mice colonized with *H. bilis* and exposed to low-dose DSS is the requirement for a microbiota. The microbiota need not be complex, as severe disease was detected in mice with a conventional community and in those harboring the simplified ASF consortium. Involvement of the microbiota in pathobiont-initiated disease has also been observed during *H. hepaticus*-driven colitis ^32,33^ as well as during the induction of disease in immunocompromised hosts by the pathobionts *P. mirabilis* and *K. pneumoniae* ^9,10,34^. Collectively, these studies and our own observations for *H. bilis* highlight the fact that multiple species of pathobiont act synergistically with resident symbionts to exacerbate pathology. It is important to note that the requirement of a microbiota for disease initiation and/or severity is not unique to pathobiont-mediated pathology. Indeed, the magnitude of intestinal inflammation in several genetically-induced immunodeficient mouse models (*IL-10*^*-/-*^, *IL-2*^*-/-*^, *TCR-alpha*^*-/-*^) ^35-39^ also depends on one or more members of the microbiota. Thus, the microbiota can play synergistic roles in disease processes that are initiated by entirely different mechanisms (pathobiont-mediated or genetic predisposition), suggesting that its involvement may be a general feature of inflammatory intestinal diseases.

One of the most fundamental ways in which the resident microbiota can contribute to chronic intestinal inflammation is via aberrant engagement of the adaptive immune system ^40^. In our model, we demonstrate that pathobiont-induced tissue damage is mediated, in part, by the adaptive immune system and characterized by Th17 responses toward gut symbionts. These observations are consistent with those made by Belkaid and colleagues, who show that microbiota-specific T cells are activated during acute gastrointestinal infection with *Toxoplasma gondii* and are functionally identical to pathogen-specific T cells, leading the authors to conclude that primary immune responses to intestinal pathogens likely occur in the context of secondary immune responses directed against the microbiota ^41,42^. The individual microbial targets of such T cell responses during disease are largely unknown, with the exception of flagellin ^39,43-48^. Much of the difficulty in dissecting contributions of pathobionts, specific T cell responses, and the microbiota during inflammatory intestinal disease lies in the complexity of the microbiota itself ^9^. We circumvented this problem by using a defined microbiota mouse model, in which *H. bilis* colonization and low-dose DSS treatment caused a disease phenotype indistinguishable from that of mice carrying a complex microbiota. With this reductionist approach, we were able to demonstrate that specific resident gut symbionts—primarily ASF 356 (*Clostridium sp.*), ASF 361 (*Lactobacillus murinus*) and ASF 502 (*Clostridium sp.*)—are the dominant targets of a Th17 response rather than the pathobiont itself. However, these Th17 targeted species are not selectively coated with IgA during disease as has been demonstrated in other models of intestinal inflammation^49^. The discordance we observed between Th17 and IgA immune reactivity may indicate that differential immune signatures could occur in IBD. Future studies will investigate how *H. bilis* instigates Th17 responses exclusively against the microbiota. Considering that some Helicobacter species were recently reported to precipitate antigen-specific T cell responses against themselves but not other bacterial taxa that expanded during disease, additional work to provide perspective regarding the diversity of immunological influence exerted by various Helicobacter species is warranted ^55^.

The ability of *H. bilis* to incite CD4^+^ T cell responses against the microbiota, but not itself, is a novel finding that may partly explain why no single microorganism or immune response against particular microbiota members have ever been associated with inflammatory bowel diseases. Even more striking is our observation that severe pathobiont-mediated intestinal inflammation does not appear to depend on any specific member of the microbiota, as removal of the immune-dominant taxa in the ASF community altered the primary targets, but did not influence disease severity. We believe this finding is not only of mechanistic significance, but also of great importance to IBD diagnostics, because it implies that testing for changes in the development of immune responses to the gut microbiota may be of far greater diagnostic value than testing for responses to a specific pathobiont or particular member of the microbiota.

## Materials and Methods

### Bacterial strains

*Helicobacter bilis* WiWa ^18,56^ was grown in broth overlaid onto Brucella agar plates. Agar consisted of Brucella broth (BD BBL™ Brucella Broth, Difco™ Becton Dickinson, Sparks, MD) and 0.125% activated charcoal (Acros Organics, Thermo Fisher Scientific Inc., Fair Lawn, NJ) adjusted to pH 5 using 6M HCl solution prior to the addition of 1.6% bacteriological agar (Amresco®, Solon, OH). After autoclaving, cooled agar was with supplemented with 20% heat-inactivated newborn calf serum (hi-NCS; HyClone®, Thermo Fisher Scientific Inc.) and 2% Vitox supplement (Oxoid™, Thermo Fisher Scientific Inc.) prior to pouring plates. The broth overlay consisted of 70% Brucella broth (autoclaved and cooled prior to use), 12% urea (BD BBL™ Urea Agar Base, Difco™ Becton Dickinson), 17% hi-NCS and 1% isovitalex supplement (BBL™ Medium Enrichment for Fastidious Microorganisms, BD). *H. bilis* cultures were maintained in modular incubator chambers (Billups-Rothenberg, Del Mar, CA) under microaerophilic conditions (80% N_2_, 10% H_2_ and 10 % CO_2_) at 37°C. The broth overlay was changed every 12 hr for 36 hr prior to harvesting *H. bilis* for inoculation. All cultures were continuously checked for aerobic contaminants via plating on Tryptic Soy Agar plates (BD Tryptic Soy Agar, Difco™ Becton Dickinson) at 37°C. Prior to inoculation, *H. bilis* cultures were evaluated for purity, morphology and motility via dark-phase microscopy. Bacteria were then suspended in Brucella broth (BD BBL™ Brucella Broth, Difco™ Becton Dickinson) at an approximate concentration of 10^8^ CFU/mL using MacFarland standards.

The ASF microbial community consisted of ASF 356, *Clostridium sp.*; ASF 360, *Lactobacillus intestinalis*; ASF 361, *Lactobacillus murinus*; ASF 457, *Mucispirillum schaedleri*; ASF 492, *Eubacterium plexicaudatum*; ASF 500, *Pseudoflavonifractor* sp.; ASF 502, *Clostridium* sp.; and ASF 519, *Parabacteroides goldsteinii* ^57,58^. Individual ASF members were cultured under anaerobic conditions as previously described ^59^. ASF strains, *H. bilis* WiWa, and founder C3H/HeN gnotobiotic breeding mice harboring all eight ASF members were a kind gift from Dr. Michael Wannemuehler, Iowa State University.

### Animal experiments

Conventional (Conv) and ASF-bearing C3H/HeN mice, ASF-bearing C57BL/6 or *Rag1*^*-/-*^ mice, germ-free (GF) C3H/HeN, and GF *Rag1*^-/-^ mice were bred and maintained at the University of Nebraska-Lincoln Gnotobiotic Mouse Facility. All GF and ASF-bearing mice were bred and reared under gnotobiotic conditions in flexible film isolators. All mice were fed an autoclaved chow diet ad libitum (LabDiet 5K67, Purina Foods). Germ-free status was routinely checked as previously described ^59^. Conventional mice were housed in autoclaved cages on a positive pressure, individually ventilated caging system (Allentown Inc., Allentown, NJ), maintained on autoclaved bedding, provided autoclaved water and fed an autoclaved diet. Cages were only opened in a biosafety cabinet. ASF-bearing mice used in experiments were colonized with the ASF via vertical transmission (i.e., from birth). ASF colonization status of breeding and experimental mice was verified using fecal samples and previously published qPCR assays ^59^. Mice inoculated with *H. bilis* received a single oral gavage containing 10^8^ CFU/mL in 200 μL. *H. bilis* colonization was verified 7-10 days post-inoculation using fecal samples and a qPCR assay. All control Conv and ASF-bearing animals were confirmed to be devoid of *H. bilis*.

Defined microbial communities consisting of various ASF members were assembled in adult germ-free C3H/HeN mice by first resuspending pure cultures of each individual ASF member to a density of 10^6^-10^8^ CFU/mL using McFarland standards. Mice received a 200 uL oral gavage of each organism daily for three consecutive days. Community assembly was achieved by first introducing ASF 360 (*L. intestinalis*) and 519 (*P. goldsteinii*); remaining ASF members were then added in the following order: ASF 500 (*Pseudoflavonifractor sp.*), 502 (*Clostridium sp.*), 492 (*E. plexicaudatum*), 356 (*Clostridium sp.*), 361 (*L. murinus*), and 457 (*M. schaedleri*).

To trigger disease, mice were provided with either 1.5% (for all C3H/HeN) or 2% (all C57BL/6 WT and *Rag1*^*-*/-^) dextran sulfate sodium salt (DSS; MW = 36,000 - 50,000, MP Biomedicals, LLC, Solon, OH) in their drinking water for 5 days, followed by a restitution period with regular water for 4 days prior to necropsy. Age-matched control animals received regular drinking water for the duration of the study. Considering previously published data^18^, each experiment measuring disease phenotype as well as immunological and bacteriological parameters consisted of at least 6-8 animals per treatment and each experiment was performed once or twice. In some experiments, sample size was adjusted to maximize statistical power as the data for most parameters measured were not normally distributed and cell samples needed to be pooled for select assays. Animals were not randomly distributed across treatments. The Institutional Animal Care and Use Committee at the University of Nebraska-Lincoln approved all procedures involving animals (Protocols 817 and 1215).

### DNA isolation and quantitative real-time PCR

DNA was isolated from feces and cecal contents as previously described using a phenol-chloroform-isoamyl alcohol and chloroform-isoamyl alcohol based protocol ^60^. Cecal tissue DNA was isolated using the QIAamp DNA Blood and Tissue Kit as per manufacturer instructions (QIAamp DNA Blood and Tissue Kit, Qiagen(®), Germantown, MD). DNA was quantified using fluorescent molecule labeling, and all samples were diluted to a final concentration of 10 ng/μL prior to using 1 μL of the DNA template in each qPCR reaction as previously described ^59^. Quantification of all ASF members in wet cecal contents and tissues was performed using previously described qPCR assays. The abundance of *H. bilis* was quantified using a newly designed primer set and qPCR assay^59^. Primer sequences were Forward: 5’ TGGCTTGCCAGAGCTTGA 3’ and Reverse: 5’ CTGCTAGCAACTAAGGACG 3’ (IDT DNA Technologies, Coralville, IA). Amplicon size was 111 base pairs. Thermocycling conditions included: (i) an initial denaturation step of 10 min at 95°C; (ii) 35 cycles of 20 s at 95°C, 30 s at 60°C (annealing temperature), and 45 s at 68°C; (iii) one cycle of 20 s at 95°C; (iv) one cycle of 30 s at 60°C; (v) one 20 min interval to generate a melting curve; and (vi) one cycle of 45 s at 95°C. Both forward and reverse primers were used at a final concentration of 300 nM. See Supplementary Information for more detailed information.

### Gross and histopathological lesion scores

At necropsy, a gross cecal disease score was assigned using a previously described set of criteria ^61^ modified to include atrophy, emptying, enlargement of the cecal tonsil, presence of mucoid contents, and presence of intraluminal blood. For each parameter, a score of 0 (absent) or 1 (present) was given. Final scores were additive for all parameters; the higher the final gross score, the greater the disease severity. The investigator assigning gross scores was not blinded to treatments; however, tissues from healthy control mice were always scored first to establish baseline values. To assess microscopic lesions, the apical portions of cecal tissues were collected, fixed in 10% neutral buffered formalin (Thermo Fisher Scientific Inc.) and subsequently processed, sectioned and stained with hematoxylin and eosin (H&E). Tissues were scored by a board-certified veterinary pathologist (J.M.H.) who was blinded to treatments. Cumulative histopathological scores ranged from 0-30 and were based on a 0 to 5 score for each of the following parameters adapted from a previous report ^61^: gland hyperplasia, stromal collapse, edema, cellular inflammation, ulceration and mucosal height. Higher cumulative scores represented more severe disease.

### Cecal explant cultures

Cultures of cecal fragments were prepared as previously described for colon fragments ^37^. More detailed information can be found in the Supplementary Material.

### CD4^+^ T cell stimulation

A culture system was adapted from a previously described protocol ^62^ as a method to provide purified CD4^+^ T cells with an *in vitro* stimulation to assess antigen-specific cytokine production in response to *H. bilis* or individual ASF antigens. See Supplementary Information for more detailed information.

### Cytokine quantification

Chemokine and cytokine levels (i.e., from cecal explants or CD4^+^ T cell supernatants) were measured using customized Mouse Cytokine/Chemokine Magnetic Bead Kits (Milliplex, Millipore, Billerica, MA) and a MAGPIX instrument (Luminex Corporation, Austin, TX), except for TGF-ß1, which was measured using a Mouse TGF-ß1 ELISA Ready-SET-Go (eBioscience Inc.) as per manufacturer instructions.

### Flow cytometry

Immune cell phenotypes were assessed as previously described ^63,64^. See Supplementary Information for more detailed information and schematics of gating strategies (Fig. S11A-E and S12A-F).

### Immunoglobulin A (IgA)-based bacterial cell sorting and quantification

This method was adapted from a previously published work^49^. A detailed description of the protocol used for bacterial cell sorting and quantification can be found in the Supplementary Material.

### Statistical analysis

Statistical analyses were performed using GraphPad Prism 7 (version 7.0a, 2016, GraphPad Software, Inc., La Jolla, CA) and the R software, version 3.3.1 (R Core Team 2016, R Foundation for Statistical Computing, Vienna, Austria). More detailed description of all analyses can be found in the Supplementary Information. Significant differences between two treatments are presented by * = *P* < 0.05, ** = *P* ≤ 0.01, *** = *P* ≤ 0.001, **** = *P* ≤ 0.0001 and NS = not significant at *P* ≥ 0.05.

### Data availability

R scripts and raw data for the ecological analysis (i.e., ASF community structure and correlation analysis) are accessible at https://github.com/jcgneto/HbilisASFmodel.

## Acknowledgments

We are extremely grateful for the technical expertise and skillful animal husbandry provided by Brandon White and the staff at the UNL Gnotobiotic Mouse Facility. We would also like to acknowledge the expert technical support provided by Dirk Anderson, UNL Center for Biotechnology Flow Cytometry Core. We especially wish to thank Dr. Michael Wannemuehler (Iowa State University) for generously providing founder ASF-bearing breeding C3H/HeN mice, ASF strains and *H. bilis* WiWa. We also thank Drs. Stephen Kachman, Jay Reddy and Deborah Brown from the University of Nebraska-Lincoln for helpful discussions. This work was supported by a Career Development Award from the Crohn’s and Colitis Foundation of America (grant # 3578), the National Institute of General Medical Sciences of the National Institutes of Health (1P20GM104320), and start-up funding from the University of Nebraska-Lincoln to A.E.R.T. L.B.B. was supported by a complementary post-doctoral grant awarded by the Fonds Spécial de Recherche, Université catholique de Louvain. The funders had no role in study design, data collection and analysis, decision to publish or preparation of the manuscript.

### Author Contributions

J.C.G.N. and A.E.R.T. conceived the project, designed the experiments and wrote the manuscript. J.C.G.N., H.K., S.M., R.R.S.M., R.J.S. and L.B.B. performed experiments. J.C. provided statistical support. J.M.H. was the veterinary pathologist who blindly scored tissues. A.K.B. and J.W. contributed to experimental design, data interpretation and manuscript preparation.

## Additional Information

### Supplementary information

accompanies this paper.

## References

1 Huttenhower, C., Kostic, A. D. & Xavier, R. J. Inflammatory bowel disease as a model for translating the microbiome. Immunity 40, 843–854, (2014).

2 Sartor, R. B. Microbial influences in inflammatory bowel diseases. Gastroenterology 134, 577–594, (2008).

3 Frank, D. N. et al. Molecular-phylogenetic characterization of microbial community imbalances in human inflammatory bowel diseases. Proc Natl Acad Sci U S A 104, 13780–13785, (2007).

4 Gevers, D. et al. The treatment-naive microbiome in new-onset Crohn's disease. Cell Host Microbe 15, 382–392, (2014).

5 Chow, J., Tang, H. & Mazmanian, S. K. Pathobionts of the gastrointestinal microbiota and inflammatory disease. Curr Opin Immunol 23, 473–480, (2011).

6 Round, J. L. & Mazmanian, S. K. The gut microbiota shapes intestinal immune responses during health and disease. Nat Rev Immunol 9, 313–323, (2009).

7 Winter, S. E. & Baumler, A. J. Why related bacterial species bloom simultaneously in the gut: principles underlying the 'Like will to like' concept. Cell Microbiol 16, 179–184, (2014).

8 Winter, S. E. & Baumler, A. J. Dysbiosis in the inflamed intestine: chance favors the prepared microbe. Gut Microbes 5, 71–73, (2014).

9 Garrett, W. S. et al. Enterobacteriaceae act in concert with the gut microbiota to induce spontaneous and maternally transmitted colitis. Cell Host Microbe 8, 292–300, (2010).

10 Garrett, W. S. et al. Communicable ulcerative colitis induced by T-bet deficiency in the innate immune system. Cell 131, 33–45, (2007).

11 Chassaing, B., Koren, O., Carvalho, F. A., Ley, R. E. & Gewirtz, A. T. AIEC pathobiont instigates chronic colitis in susceptible hosts by altering microbiota composition. Gut 63, 1069–1080, (2014).

12 Elinav, E. et al. NLRP6 inflammasome regulates colonic microbial ecology and risk for colitis. Cell 145, 745–757, (2011).

13 Seo, S. U. et al. Distinct commensals induce interleukin-1beta via NLRP3 inflammasome in inflammatory monocytes to promote intestinal inflammation in response to injury. Immunity 42, 744–755, (2015).

14 Devkota, S. et al. Dietary-fat-induced taurocholic acid promotes pathobiont expansion and colitis in Il10-/-mice. Nature 487, 104–108, (2012).

15 Zhang, Q. et al. Accelerated dysbiosis of gut microbiota during aggravation of DSS-induced colitis by a butyrate-producing bacterium. Sci Rep 6, 27572, (2016).

16 Bloom, S. M. et al. Commensal Bacteroides species induce colitis in host-genotype-specific fashion in a mouse model of inflammatory bowel disease. Cell Host Microbe 9, 390–403, (2011).

17 Chow, J. & Mazmanian, S. K. A pathobiont of the microbiota balances host colonization and intestinal inflammation. Cell Host Microbe 7, 265–276, (2010).

18 Liu, Z. et al. Helicobacter bilis colonization enhances susceptibility to typhlocolitis following an inflammatory trigger. Dig Dis Sci 56, 2838–2848, (2011).

19 Fox, J. G. et al. *Helicobacter bilis* sp. nov., a novel Helicobacter species isolated from bile, livers, and intestines of aged, inbred mice. J Clin Microbiol 33, 445–454, (1995).

20 Maggio-Price, L. et al. Dual infection with *Helicobacter bilis* and *Helicobacter hepaticus* in p-glycoprotein-deficient mdr1a-/-mice results in colitis that progresses to dysplasia. Am J Pathol 166, 1793–1806, (2005).

21 Maggio-Price, L. et al. *Helicobacter bilis* infection accelerates and *H. hepaticus* infection delays the development of colitis in multiple drug resistance-deficient (mdr1a-/-) mice. Am J Pathol 160, 739–751, (2002).

22 Maxwell, J. R. et al. Differential roles for interleukin-23 and interleukin-17 in intestinal immunoregulation. Immunity 43, 739–750, (2015).

23 Khor, B., Gardet, A. & Xavier, R. J. Genetics and pathogenesis of inflammatory bowel disease. Nature 474, 307–317, (2011).

24 Sartor, R. B. Mechanisms of disease: pathogenesis of Crohn’s disease and ulcerative colitis. Nat Clin Pract Gastroenterol Hepatol 3, 390–407, (2006).

25 Maloy, K. J. & Powrie, F. Intestinal homeostasis and its breakdown in inflammatory bowel disease. Nature 474, 298–306, (2011).

26 Lupp, C. et al. Host-mediated inflammation disrupts the intestinal microbiota and promotes the overgrowth of Enterobacteriaceae. Cell Host Microbe 2, 204, (2007).

27 Papa, E. et al. Non-invasive mapping of the gastrointestinal microbiota identifies children with inflammatory bowel disease. PLoS One 7, e39242, (2012).

28 Mukhopadhya, I., Hansen, R., El-Omar, E. M. & Hold, G. L. IBD-what role do Proteobacteria play? Nat Rev Gastroenterol Hepatol 9, 219–230, (2012).

29 Haines, D. C. et al. Inflammatory large bowel disease in immunodeficient rats naturally and experimentally infected with *Helicobacter bilis*. Vet Pathol 35, 202–208, (1998).

30 Carvalho, F. A. et al. Transient inability to manage proteobacteria promotes chronic gut inflammation in TLR5-deficient mice. Cell Host Microbe 12, 139–152, (2012).

31 Atherly, T. et al. *Helicobacter bilis* infection alters mucosal bacteria and modulates colitis development in defined microbiota mice. Inflamm Bowel Dis 22, 2571–2581, (2016).

32 Yang, I. et al. Intestinal microbiota composition of interleukin-10 deficient C57BL/6J mice and susceptibility to *Helicobacter hepaticus*-induced colitis. PLoS One 8, e70783, (2013).

33 Nagalingam, N. A. et al. The effects of intestinal microbial community structure on disease manifestation in IL-10-/-mice infected with *Helicobacter hepaticus*. Microbiome 1, 15, (2013).

34 Dieleman, L. A. et al. *Helicobacter hepaticus* does not induce or potentiate colitis in interleukin-10-deficient mice. Infect Immun 68, 5107–5113, (2000).

35 Contractor, N. V. et al. Lymphoid hyperplasia, autoimmunity, and compromised intestinal intraepithelial lymphocyte development in colitis-free gnotobiotic IL-2-deficient mice. J Immunol 160, 385–394, (1998).

36 Dianda, L. et al. T cell receptor-alpha beta-deficient mice fail to develop colitis in the absence of a microbial environment. Am J Pathol 150, 91–97, (1997).

37 Sellon, R. K. et al. Resident enteric bacteria are necessary for development of spontaneous colitis and immune system activation in interleukin-10-deficient mice. Infect Immun 66, 5224–5231, (1998).

38 Kim, S. C., Tonkonogy, S. L., Karrasch, T., Jobin, C. & Sartor, R. B. Dual-association of gnotobiotic IL-10-/-mice with 2 nonpathogenic commensal bacteria induces aggressive pancolitis. Inflamm Bowel Dis 13, 1457–1466, (2007).

39 Cong, Y. et al. CD4+ T cells reactive to enteric bacterial antigens in spontaneously colitic C3H/HeJBir mice: increased T helper cell type 1 response and ability to transfer disease. J Exp Med 187, 855–864, (1998).

40 Littman, D. R. & Rudensky, A. Y. Th17 and regulatory T cells in mediating and restraining inflammation. Cell 140, 845–858, (2010).

41 Hand, T. W. et al. Acute gastrointestinal infection induces long-lived microbiota-specific T cell responses. Science 337, 1553–1556, (2012).

42 Belkaid, Y. & Hand, T. W. Role of the microbiota in immunity and inflammation. Cell 157, 121–141, (2014).

43 Cong, Y., Weaver, C. T., Lazenby, A. & Elson, C. O. Colitis induced by enteric bacterial antigen-specific CD4+ T cells requires CD40-CD40 ligand interactions for a sustained increase in mucosal IL-12. J Immunol 165, 2173–2182, (2000).

44 Lodes, M. J. et al. Bacterial flagellin is a dominant antigen in Crohn disease. J Clin Invest 113, 1296–1306, (2004).

45 Brimnes, J., Reimann, J., Nissen, M. & Claesson, M. Enteric bacterial antigens activate CD4(+) T cells from scid mice with inflammatory bowel disease. Eur J Immunol 31, 23–31, (2001).

46 Ivanov, II et al. Induction of intestinal Th17 cells by segmented filamentous bacteria. Cell 139, 485–498, (2009).

47 Eun, C. S. et al. Induction of bacterial antigen-specific colitis by a simplified human microbiota consortium in gnotobiotic interleukin-10-/-mice. Infect Immun 82, 2239–2246, (2014).

48 Kullberg, M. C. et al. Induction of colitis by a CD4+ T cell clone specific for a bacterial epitope. Proc Natl Acad Sci U S A 100, 15830–15835, (2003).

49 Palm, N. W. et al. Immunoglobulin A coating identifies colitogenic bacteria in inflammatory bowel disease. Cell 158, 1000–1010, (2014).

50 Kullberg, M. C. et al. IL-23 plays a key role in *Helicobacter hepaticus*-induced T cell-dependent colitis. J Exp Med 203, 2485–2494, (2006).

51 Hue, S. et al. Interleukin-23 drives innate and T cell-mediated intestinal inflammation. J Exp Med 203, 2473–2483, (2006).

52 Leppkes, M. et al. RORgamma-expressing Th17 cells induce murine chronic intestinal inflammation via redundant effects of IL-17A and IL-17F. Gastroenterology 136, 257–267, (2009).

53 Rivas, M. A. et al. Deep resequencing of GWAS loci identifies independent rare variants associated with inflammatory bowel disease. Nat Genet 43, 1066–1073, (2011).

54 Barrett, J. C. et al. Genome-wide association defines more than 30 distinct susceptibility loci for Crohn’s disease. Nat Genet 40, 955–962, (2008).

55 Chai, J. N. et al. Helicobacter species are potent drivers of colonic T cell responses in homeostasis and inflammation. Sci Immunol 2, (2017).

56 Jergens, A. E. et al. *Helicobacter bilis* triggers persistent immune reactivity to antigens derived from the commensal bacteria in gnotobiotic C3H/HeN mice. Gut 56, 934–940, (2007).

57 Dewhirst, F. E. et al. Phylogeny of the defined murine microbiota: altered Schaedler flora. Appl Environ Microbiol 65, 3287–3292, (1999).

58 Wymore Brand, M. et al. The Altered Schaedler Flora: Continued applications of a defined murine microbial community. ILAR J 56, 169–178, (2015).

59 Gomes-Neto, J. C. et al. A real-time PCR assay for accurate quantification of the individual members of the Altered Schaedler Flora microbiota in gnotobiotic mice. J Microbiol Methods 135, 52–62, (2017).

60 Martinez, I. et al. Diet-induced metabolic improvements in a hamster model of hypercholesterolemia are strongly linked to alterations of the gut microbiota. Appl Environ Microbiol 75, 4175–4184, (2009).

61 Jergens, A. E. et al. Induction of differential immune reactivity to members of the flora of gnotobiotic mice following colonization with *Helicobacter bilis* or *Brachyspira hyodysenteriae*. Microbes Infect 8, 1602–1610, (2006).

62 Ramer-Tait, A. E., Petersen, C. A. & Jones, D. E. IL-2 limits IL-12 enhanced lymphocyte proliferation during *Leishmania amazonensis* infection. Cell Immunol 270, 32–39, (2011).

63 Ramer, A. E., Vanloubbeeck, Y. F. & Jones, D. E. Antigen-responsive CD4+ T cells from C3H mice chronically infected with *Leishmania amazonensis* are impaired in the transition to an effector phenotype. Infect Immun 74, 1547–1554, (2006).

64 Jones, D. E., Buxbaum, L. U. & Scott, P. IL-4-independent inhibition of IL-12 responsiveness during *Leishmania amazonensis* infection. J Immunol 165, 364–372, (2000).

